# Pan-cancer Analysis Identified Ectopic RUNX1T1 Associated with Lineage Plasticity

**DOI:** 10.1101/2025.04.18.649575

**Authors:** Yuyin Jiang, Siyuan Cheng, Isaac Yi Kim, Su Deng, Ping Mu

**Affiliations:** Department of Urology, Yale University School of Medicine, New Haven, CT, 06511; Yale Cancer Center, Yale University School of Medicine, New Haven, CT, 06511

**Author notes:** Correspondence to (P.M.), (S.D.). These authors contributed equally to this work (Yuyin Jiang, Siyuan Cheng).

**Keywords:** HOX code, Lineage plasticity, Prostate cancer, Lung cancer, AML, *RUNX1T1*

## Abstract

Cancer remains a leading cause of death worldwide, with lineage plasticity emerging as a hallmark that drives therapy resistance and tumor progression, allowing cancer cells to rapidly alter their identity and evade targeted therapies. Although various genomic and transcriptomic aberrations correlate with lineage plasticity, the lack of pan-cancer gene markers quantifying lineage plasticity has limited its utility as a predictive biomarker. Homeobox (*HOX*) genes encode a family of transcription factors that play critical roles in embryonic development and tissue identity by establishing distinct expression patterns, known as HOX codes, in specific cell lineages. Through comprehensive bioinformatic analysis of multi-omics data—including expression profiles of 39 *HOX* genes from over 80,000 RNA sequencing samples across 114 cancer types—we first demonstrated that HOX codes effectively represent the lineages of cancer cells. We then identified multiple lineage-plastic cancer subtypes by applying the calculated HOX codes. Specifically, we identified lineage-plastic tumor subtypes in prostate cancer, lung cancer, and acute myeloid leukemia (AML), which exhibit altered HOX codes compared to non-plastic subtypes. Downstream differential expression analysis revealed significantly elevated *RUNX1T1* levels across all three lineage-plastic cancer types, which was further validated through bulk and single-cell RNA sequencing data derived from preclinical and clinical samples. Together, our findings provide a novel strategy for characterizing lineage plasticity in pan-cancer cells and suggest ectopic *RUNX1T1* expression as a pan-cancer marker and critical mediator of lineage plasticity.

## Introduction

Cancer remains the second leading cause of death worldwide, and despite advances in treatment, most cancers eventually develop drug resistance and metastasize. Cancer cells possess the ability to alter their established lineage by reverting to a stem-like state and subsequently re-differentiating into an alternative lineage, thereby evading therapies targeting their original lineage-directed survival programs [1-4]. This plasticity has been demonstrated to cause resistance to many standard cancer therapies in various types of human cancers, including prostate, breast, lung, and pancreatic cancers, as well as melanoma [2, 4-10]. Prostate cancer (PCa) represents a salient example of how cancer cells acquire resistance through lineage plasticity. PCa is the second most frequently diagnosed cancer among American men, with the deadliest form being metastatic castration-resistant prostate cancer (mCRPC), which is currently incurable and is responsible for almost all PCa related death [11, 12]. Although the development of second-generation AR-targeted therapies has significantly improved the survival of men with mCRPC, resistance to these agents is unfortunately inevitable, largely limiting the clinical outcomes of patients with mCRPC [13-15]. Emerging evidence demonstrates that the luminal type of mCRPC can transform into lineage-plastic types of cancers, including neuroendocrine prostate cancer (NEPC), double-negative mCRPC, or progenitor-type mCRPC, which become independent of androgens and highly resistant to therapies. Similarly, in lung cancer, lineage plasticity contributes to the transformation of lung adenocarcinoma (LUAD) to small cell lung cancer (SCLC) or squamous cell carcinoma, resulting in resistance against EGFR inhibitors. Although less studied, lineage plasticity has been reported in other cancer types such as bladder, pancreas, esophageal cancer, and glioblastoma [16].

Efforts to decode the mechanisms driving lineage plasticity have revealed key contributors in specific cancers. For example, genetic loss of *TP53* and *RB1* has been shown to facilitate lineage plasticity in prostate cancer and lung cancer [4, 17]. Across different cancer types, epigenetic regulators such as Tet Methylcytosine Dioxygenase 2 (TET2) [18] and Enhancer of Zeste 2 Polycomb Repressive Complex 2 Subunit (EZH2) [19], along with transcription factors including SRY-Box Transcription Factor 2 (SOX2) [4] and Achaete-Scute Family BHLH Transcription Factor 1 (ASCL1) [20, 21], have been reported to contribute to lineage plasticity and targeted therapy resistance in various cancers. Other epigenetic modifiers such as CHD1, LSD1, and the SWI/SNF complex [5, 7, 22-27], also contribute to the acquisition of lineage plasticity. Additionally, signaling pathways such as JAK-STAT [2], RNA-binding proteins like SYNCRIP [28], and ubiquitination-related enzymes such as UBE2J1 [29] have been implicated in regulating lineage plasticity and therapy resistance in mCRPC. Despite these recent advances revealing key regulators of lineage plasticity, a pan-cancer gene marker quantifying lineage plasticity is still missing, which limits the definition and quantification of lineage plasticity across diverse cancer types and lineages. This gap in knowledge largely limits the clinical utilization of lineage plasticity as a predictive biomarker for targeted therapy resistance and hinders the practice of precision oncology.

The HOX code, defined by the spatially and temporally coordinated expression of Homeobox (*HOX*) genes, serves as a critical determinant of embryonic patterning and cell lineage identity during the normal developmental process [30]. In humans, the 39 *HOX* genes are organized into four clusters (*HOXA–D*), with their genomic arrangement mirroring their sequential activation during development: 3’-located genes are expressed earlier in anterior regions, while 5’-located genes are sequentially activated later in posterior regions [31]. This orchestrated expression governs anterior–posterior body segmentation and establishes tissue-specific identity, underscoring the role of the HOX code as a molecular blueprint for cellular lineage (Figure 1A) and indicating the potential link between the HOX code and lineage plasticity in PCa. Previously, we reported that lineage-plastic NEPC cells exhibit an altered HOX code compared to non-plastic PCa adenocarcinoma cells, as revealed through analysis of the Cancer Cell Line Encyclopedia (CCLE) database [32]. However, the initial exploration of the HOX code across pan-cancer types was limited due to insufficient sample size. In this study, through comprehensive bioinformatic analysis of multi-omics data—including expression profiles of 39 *HOX* genes from over 80,000 RNA sequencing samples across 114 cancer types—we first demonstrated that HOX codes effectively represent the lineages of cancer cells. We then identified lineage-plastic tumor subtypes in prostate cancer, lung cancer, and acute myeloid leukemia (AML), which exhibit altered HOX codes compared to non-plastic subtypes. More excitingly, our analysis revealed significantly elevated *RUNX1T1* levels across all three lineage-plastic cancer types, which was further validated through bulk and single-cell RNA sequencing data derived from preclinical and clinical samples. Together, our findings provide a novel strategy for characterizing lineage plasticity in pan-cancer cells through the HOX code and further suggest ectopic *RUNX1T1* expression as a pan-cancer marker and critical mediator of lineage plasticity, thereby filling the gap in utilizing lineage plasticity as a predictive biomarker for targeted therapy resistance.

**Figure 1.**
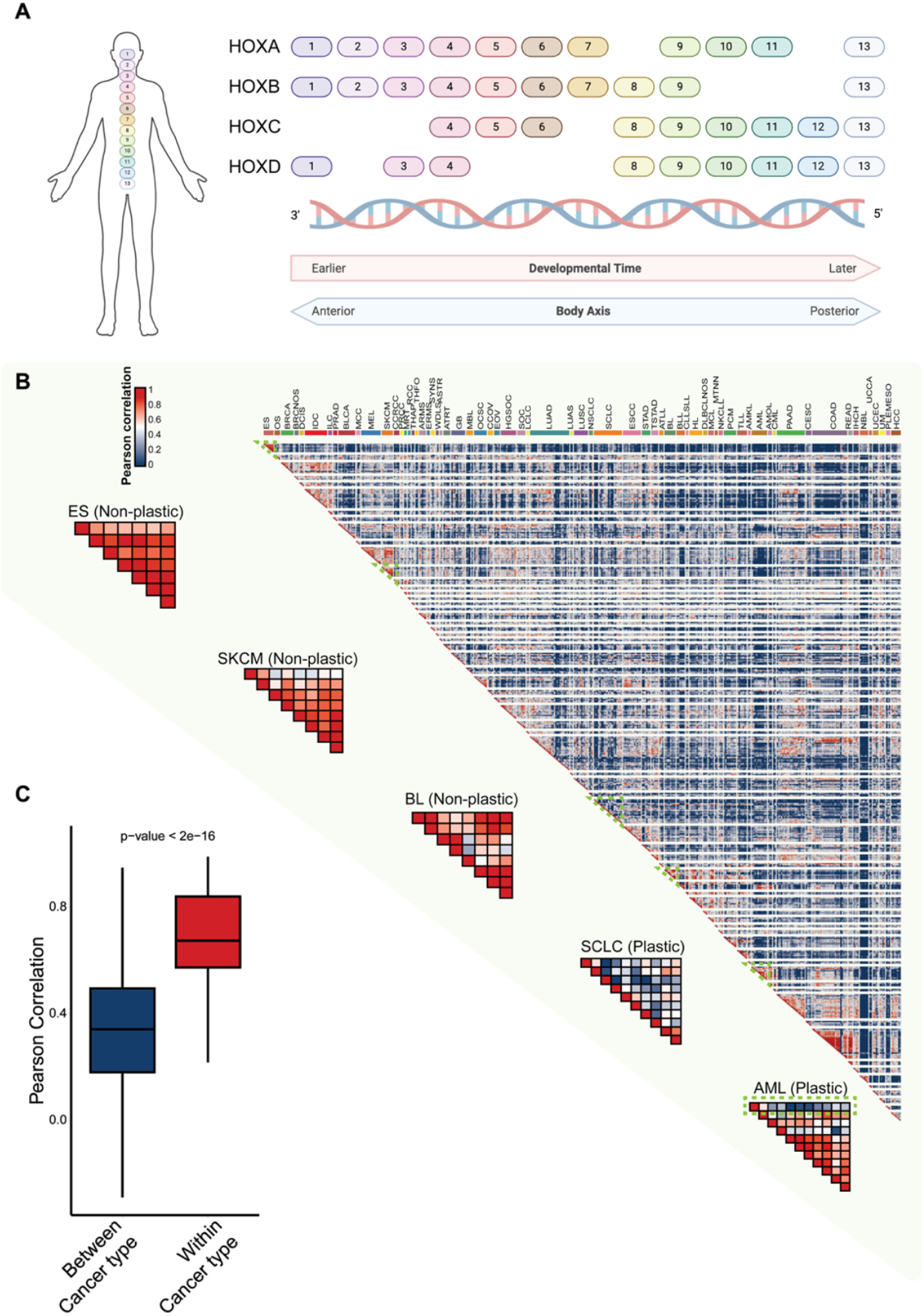
Hox code represents tissue identity and cancer cell lineages. (A) Schematic representation of *HOX* gene clusters (*HOXA-D*) arranged by embryonic developmental time and the anterior-posterior body axis. Figure was adapted from [39] and made by BioRender. (B) Heatmap showing the correlation scores between the *HOX* gene expression across cancer cell lines, ordered by cancer types. Specific portions of the heatmap representing Ewing sarcoma (ES), cutaneous melanoma (SKCM), and Burkitt lymphoma (BL), small cell lung cancer (SCLC), and acute myeloid leukemia (AML) are outlined in green and zoomed in for better visualization and comparison. (C) Average Pearson correlation test scores between and within cancer types. Difference between the two conditions is significant with a p-value of 2e-16.

## Results

### Hox Codes Represent Cancer Lineage Identity

To study whether *HOX* genes can serve as markers of cancer cell lineage plasticity, we first curated the established pan-cancer cell line database, PCTA (<u>P</u>an-cancer <u>C</u>ell Line <u>T</u>ranscriptome <u>A</u>tlas), which integrates high-quality RNA-seq data from over 80,000 RNA-seq samples across 535 cancer cell lines [33, 34]. Using these RNA-seq samples, we generated HOX codes unique to each cancer cell line, defined as the median expression of 39 *HOX* genes from all RNA-seq samples of the same cancer cell line (Table. 1). To evaluate the conservation of HOX codes within and across cancer subtypes, we performed pairwise Pearson correlation analyses and visualized the results as a heatmap, ordering cell lines by cancer type. As we expected, most cancer types exhibited robust intra-cancer type HOX code conservation, with Ewing sarcoma (ES), cutaneous melanoma (SKCM), and Burkitt lymphoma (BL) displaying particularly strong coherence (Figure 1B). For a more quantitative assessment, we grouped the HOX code correlation results based on whether the different cancer cell lines belonged to the same cancer type or different types. We found that HOX code correlations were significantly higher within the same cancer type compared to those between different cancer types (Figure 1C). These findings suggest that HOX codes may serve as markers of cancer lineage identity, extending their role from developmental biology to oncology and indicating that cancer cells retain the *HOX* expression patterns of their tissue of origin.

**Table 1.** List of 39 human HOX genes.

| Cluster | Genes | Location |
| --- | --- | --- |
| **HOXA** (11 genes) | HOXA1, HOXA2, HOXA3, HOXA4, HOXA5, HOXA6, HOXA7, HOXA9, HOXA10, HOXA11, HOXA13 | 7p15 |
| **HOXB** (10 genes) | HOXB1, HOXB2, HOXB3, HOXB4, HOXB5, HOXB6, HOXB7, HOXB8, HOXB9, HOXB13 | 17q21 |
| **HOXC** (9 genes) | HOXC4, HOXC5, HOXC6, HOXC8, HOXC9, HOXC10, HOXC11, HOXC12, HOXC13 | 12q13 |
| **HOXD** (9 genes) | HOXD1, HOXD3, HOXD4, HOXD8, HOXD9, HOXD10, HOXD11, HOXD12, HOXD13 | 2q31 |

### HOX Codes Identified Lineage-Plastic Subtypes in Prostate Cancer, Lung Cancer, and AML

To utilize HOX codes for identifying lineage plastic cancer subtypes, we first defined lineage plastic cancer cell lines as those exhibiting distinct HOX code compared to other cell lines from the same cancer type or anatomic origin and then identified two distinct patterns of lineage plasticity. Acute myeloid leukemia (AML) exemplified the first pattern in which lineage-plastic subtypes were rare, with only a small fraction of cell lines deviating from the conserved HOX code observed in most AML models (Figure 1B). The second group, represented by small cell lung cancer (SCLC), exhibited pervasive HOX code heterogeneity, with most cell lines displaying aberrant lineage transcriptional profiles (Figure 1B).

However, we recognize that some cancer types may arise from the same anatomical site, a detail not fully captured in our initial analysis. To address this, we extended our investigation by analyzing HOX codes across different sites of cancer origin. Specifically, we generated HOX codes specific to each cancer type and evaluated correlation between cancer types by comparing their unique HOX codes. Not surprisingly, this analysis revealed that most anatomic sites harbored cancer types with conserved intra-site HOX codes, likely due to the shared developmental origins of those cancers (Figure 2A). On the other hand, cancers that develop in different organs exhibited distinct HOX codes, suggesting they arise from different original cell types. For example, breast neoplasms (BNNOS) and kidney malignant rhabdoid tumors (MRT) showed completely different HOX patterns because they develop from distinct precursor cells [35, 36]. Applying our previous definition of lineage plasticity, we were able to identify a third lineage-plastic pattern in prostate cancer. Specifically, two types of prostate cancers - prostate adenocarcinomas (PRAD) and prostate small cell carcinomas (PRSCC) - displayed completely different HOX codes even though they both originate in the prostate. This difference likely occurs because PRSCC develops from PRAD through a process called trans-differentiation, where cancer cells switch their identity (lineage plasticity) [4]. Collectively, these findings suggest that HOX codes could serve as markers that represent lineage identity, while changes in these markers indicate when cancer cells switch identities and become lineage plastic.

**Figure 2.**
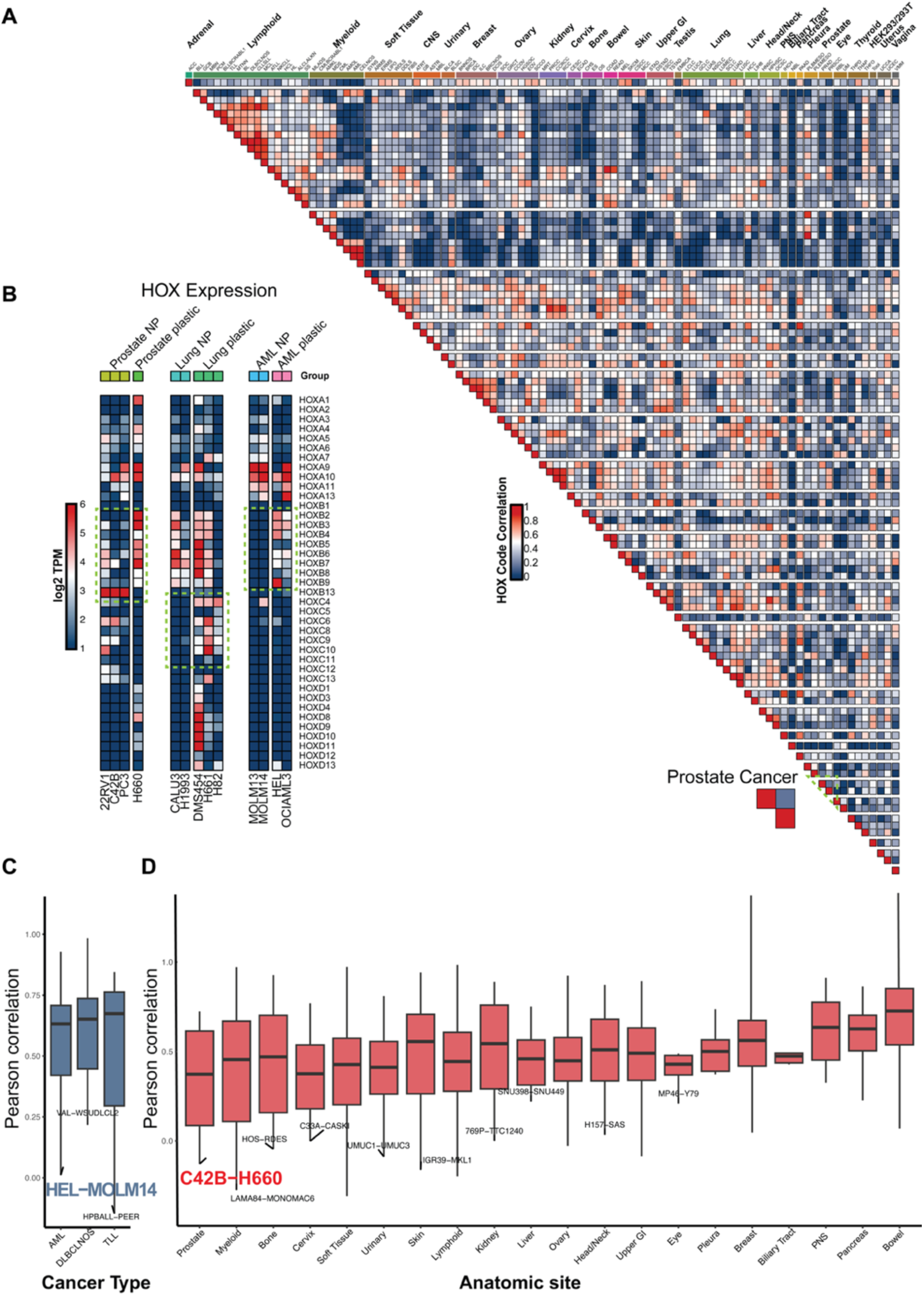
HOX codes identified lineage-plastic subtypes from prostate cancer, lung cancer, and AML. (A) Heatmap showing the correlation scores between the *HOX* gene expression across cancer types, ordered by anatomic sites. Heatmap representing prostate cancer is outlined in green and zoomed in. (B) Heatmap of *HOX* gene expression across prostate cancer, lung cancer, and AML cell lines separated in three clusters. The green box highlights distinct *HOX* gene expression patterns in lineage-plastic vs. non-plastic cell lines. NP (non-plastic) (C) Comparison of intra-cancer correlation scores across cancer types. The correlation between the HEL and MOLM14 cancer cell lines of AML is highlighted. (D) Comparison of intra-cancer correlation scores across anatomic sites. The correlation between the C4-2B and H660 prostate cancer cell lines is highlighted.

### HOX Codes Identify Preclinical Models for Dissecting Pan-Cancer Lineage Plasticity

To further dissect the lineage-plastic subtypes in prostate cancer, lung cancer, and acute myeloid leukemia (AML), we examined representative cancer cell lines using HOX codes. We visualized the HOX code correlation between cell lines through boxplots and highlighted cell line pairs with the lowest correlation across cancer types or anatomic sites (Figure 2C & 2D). Interestingly, the HOX code correlation between the lineage-plastic (HEL) and non-plastic (MOLM14) AML cell lines was the lowest compared to other pairs within the AML cancer type (Figure 2C). Similarly, across anatomical sites, the HOX codes of the prostate plastic H660 cell line and non-plastic C4-2B cell line exhibited the most significant dissimilarity (Figure 2D). These findings highlight the H660 cell line in prostate cancer and the HEL cell line in leukemia as important examples of lineage-plastic models, with HOX codes serving as clear markers of lineage plasticity differences. In contrast, we observed much higher variation between cell lines in lung cancer, which limited the visualization of these results. Therefore, we selected three lung cancer cell lines with distinct HOX codes—DMS454, H661, and H82—for further analysis (Figure 2B). To examine the differences and dynamic changes in HOX codes, we created a visual representation showing *HOX* gene expression patterns in prostate cancer, lung cancer, and AML cell lines (Figure 2B). The green box highlights prominent expression differences between lineage-plastic and non-plastic cell lines. In prostate cancer specifically, the plastic cell line exhibited elevated expression of anterior *HOX* genes (*HOXB2/3*) and reduced expression of posterior *HOX* genes (e.g., *HOXB13*) compared to non-plastic cell lines. A similar anterior-posterior HOX code shift was observed in AML (Figure 2B). Although lung cancer showed more variation between lineage-plastic cell lines, it also shared the similar pattern of reduced anterior and increased posterior *HOX* gene expression compared to non-plastic cell lines (Figure 2B).

### RUNX1T1 Is Consistently Elevated Across Lineage-Plastic Cell Lines and Cancer Types

Our analysis of HOX code patterns revealed that prostate cancer, lung cancer, and acute myeloid leukemia (AML) exhibit clear lineage plasticity, which can be quantified using HOX codes. To uncover genes associated with the acquisition of lineage plasticity across these cancer types, we retrieved all samples derived from the relevant cell lines and performed differential gene expression analysis. This analysis compared the expression profiles of over 53,000 genes between lineage-plastic and non-plastic cancers and revealed distinct transcriptional signatures associated with plasticity (Figure 3A–C). A Venn diagram identified shared gene expression patterns across the three cancer types, revealing 269 significantly upregulated genes in lineage-plastic prostate cancer, lung cancer, and AML (Figure 3D). Notably, *RUNX1T1* emerged as a top candidate gene significantly upregulated in all three lineage-plastic cancer types. To validate this finding, we generated a bar plot comparing *RUNX1T1* expression levels across plastic and non-plastic cell lines (Figure 3E). We found that elevated *RUNX1T1* expression was consistently observed across lineage-plastic cell lines in prostate cancer, lung cancer, and AML, suggesting a potential role for *RUNX1T1* in mediating pan-cancer lineage plasticity.

**Figure 3.**
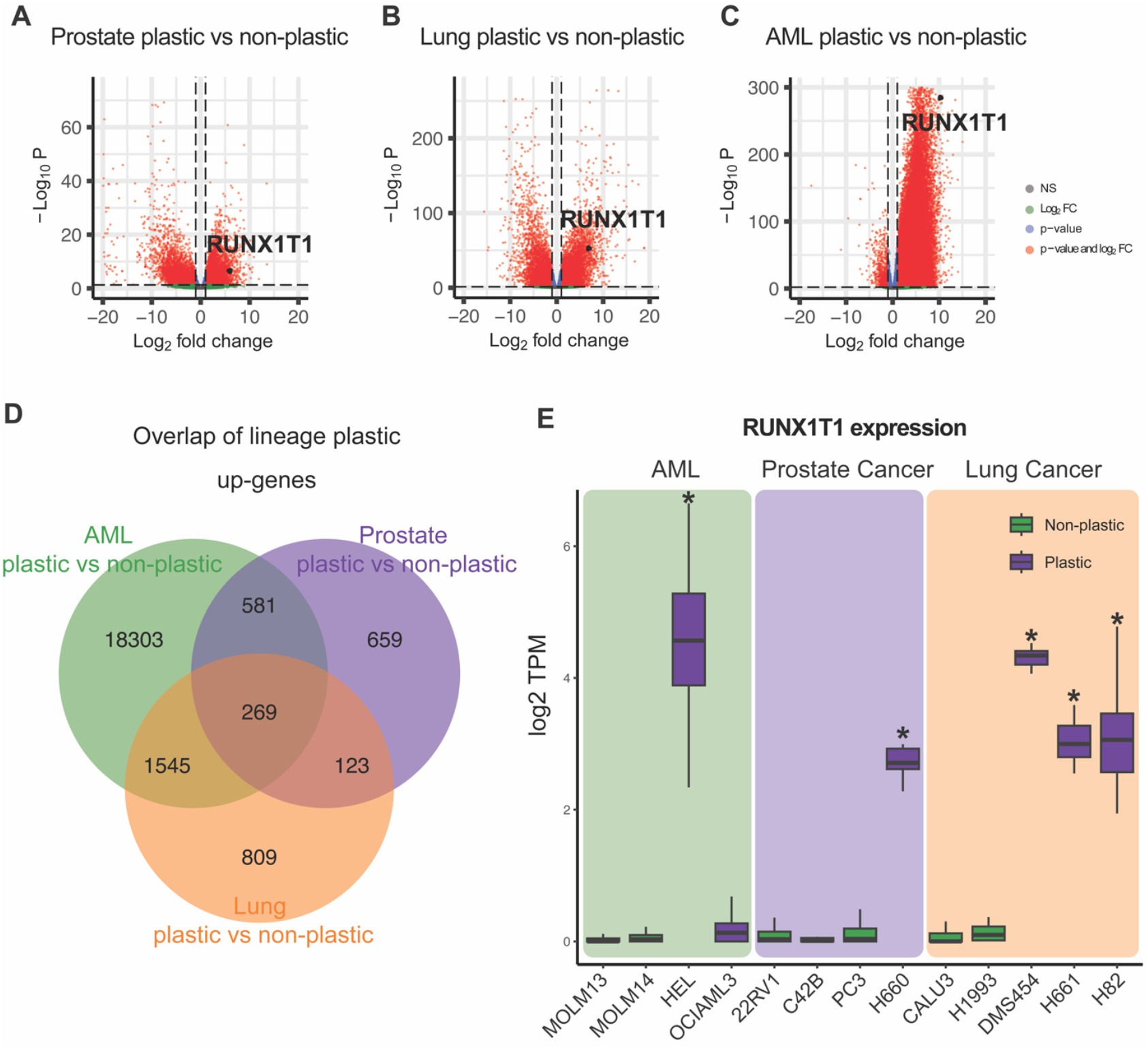
Elevated expression of RUNX1T1 in lineage plastic cell lines. (A-C) Volcano plots comparing gene expression profiles between lineage-plastic and non-plastic cancers in prostate cancer, lung cancer, and AML. A total of 53175 genes were compared. *RUNX1T1* is significantly upregulated in all three lineage-plastic cancer types. (D) Venn diagram showing 269 significantly upregulated genes shared among plastic prostate cancer, lung cancer, and AML. (E) Bar plot comparing *RUNX1T1* expression levels across lineage-plastic and non-plastic cell lines.

### ScRNA-seq and Patients Dataset Validate the Correlation of RUNXT1 with Lineage Plasticity

Building on the association of *RUNX1T1* with lineage plasticity, we next examined its expression in human and mouse single-cell models using single-cell RNA sequencing (scRNA-seq) to determine whether the findings at the bulk level were recapitulated. UMAP plots revealed distinct *RUNX1T1* expression patterns between lineage-plastic and non-plastic cancers (Figure 4A & 4E). Specifically, the lineage-plastic neuroendocrine prostate cancer showed higher *RUNX1T1* expression, while the non-plastic adenocarcinoma exhibited lower expression in both human and mouse models (Figure 4B, 4D, 4F & 4G). Consistently, UMAP analysis of *HOXB* gene expression in the human single-cell model corroborated the bulk-level HOX code patterns. The non-plastic PRAD cluster showed high expression of *HOXB13*, a marker of prostatic identity, while the plastic PRSCC cluster lost *HOXB13* expression and instead upregulated *HOXB3* and *HOXB5* (Figure 4C). Together, these results suggest that elevated *RUNX1T1* expression represents a consistent signature of lineage plasticity across various cell lines, and in both mouse and human single-cell datasets.

**Figure 4.**
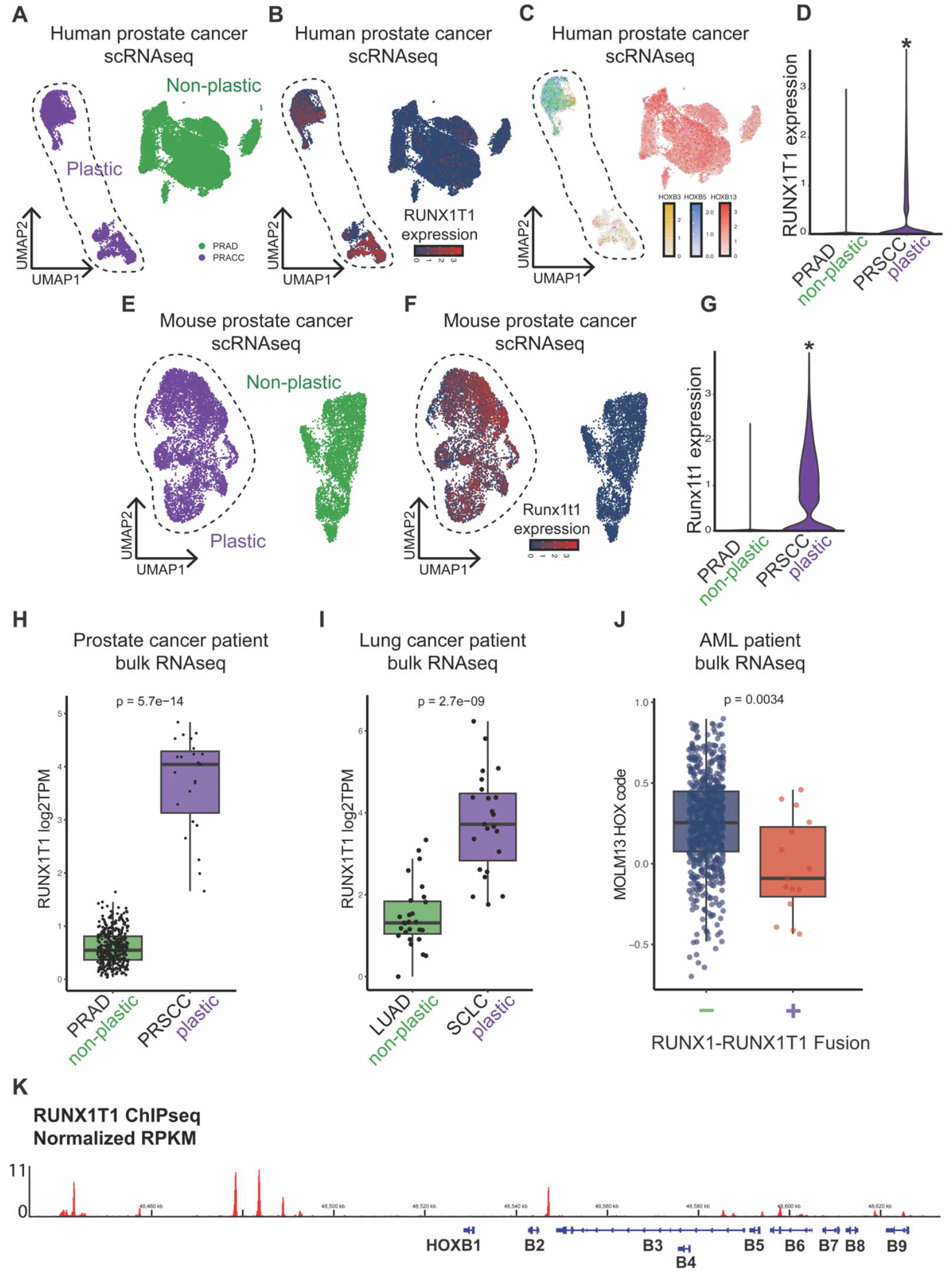
Differential expression of RUNX1T1 between lineage plastic and non-plastic human, mouse, and patient samples. (A) UMAP plots from single-cell RNA sequencing (scRNA-seq) analysis of human single-cell model. Green cluster represents non-plastic adenocarcinoma prostate cancer and purple cluster represents plastic neuroendocrine prostate cancer. (B) UMAP plot of *RUNX1T1* expression of human single-cell model showing higher *RUNX1T1* expression in lineage plastic cluster. (C) UMAP analysis of *HOXB* gene expression of human single-cell model showing loss of *HOXB13* and upregulation of *HOXB3* and *HOXB5* in lineage plastic prostate cancer. (D) Violin plots highlighting greater *RUNX1T1* expression and variability in PRSCC of human single-cell model. (E) UMAP plots from scRNA-seq analysis of mouse single-cell models. (F) UMAP plot of *RUNX1T1* expression of mouse single-cell model. (G) Violin plots of *RUNX1T1* expression in mouse PRAD and PRSCC. (H) Comparison of *RUNX1T1* expression between lineage plastic PRSCC and non-plastic PRAD patient samples in prostate cancers. (I) Comparison of *RUNX1T1* expression between lineage plastic SCLC and non-plastic LUAD patient samples in lung cancers. (J) Pearson correlation analysis of RUNX1-RUNX1T1-fused (+) and non-RUNX1-RUNX1T1-fused (−) AML samples against the non-plastic MOLM13 HOX code, showing reduced correlation in RUNX1T1-fused samples. (K) ChIP-sequencing result of the RUNX1T1 protein.

A similar trend was observed in bulk RNA-seq data from patient samples of prostate cancer, lung cancer, and AML. We compared *RUNX1T1* expression between lineage-plastic and non-plastic patient samples in prostate and lung cancers (Figure 4H & 4I). In both cancer types, the lineage-plastic subtypes—PRSCC in prostate cancer and SCLC in lung cancer—exhibited significantly higher *RUNX1T1* expression compared to their non-plastic counterparts (PRAD and LUAD, respectively). In AML, interpretation of *RUNX1T1* expression is complicated by the presence of the RUNX1–RUNX1T1 fusion. To resolve this, we separated patient samples into two groups—those with the fusion and those without—and compared their HOX code similarity to the non-plastic MOLM13 reference cell line. This analysis tested whether the association between *RUNX1T1* and plasticity persisted despite the fusion’s confounding effects. As expected, patient samples without the fusion showed higher correlation with the non-plastic HOX code, while those with the fusion exhibited significantly lower correlation (Figure 4J), indicating a loss of non-plastic characteristics due to *RUNX1T1* involvement. This finding aligns with our observations in prostate and lung cancers, reinforcing *RUNX1T1* as a pan-cancer marker of lineage plasticity across multiple cancer types. Finally, visualization of publicly available RUNX1T1 ChIP-sequencing data identified binding of the RUNX1T1 protein at the human *HOXB* genomic locus (Figure 4K), suggesting its direct regulatory function on *HOX* gene expression changes observed throughout this study.

## Discussion

Emerging evidence has highlighted the key role of lineage plasticity in driving resistance to many standard cancer therapies across various types of human cancers, including prostate, breast, lung, and pancreatic cancers, as well as melanoma [2, 4-10]. Many known factors driving lineage plasticity have been identified in different cancer types, such as loss of *TP53* and *RB1* [4, 17], ectopic activation of SOX2, TET2, EZH2, JAK-STAT, and ASCL1 [2, 4, 18, 20, 21], and modification of epigenetic regulators such as CHD1, LSD1, and the SWI/SNF complex [5, 7, 22-27]. Despite these recent advances revealing key regulators of lineage plasticity, a pan-cancer gene marker that quantifies lineage plasticity is still lacking. This absence limits the definition and quantification of lineage plasticity across diverse cancer types and lineages. This knowledge gap significantly restricts the clinical utilization of lineage plasticity as a predictive biomarker for targeted therapy resistance and hinders the practice of precision oncology. Our study demonstrates that the role of the HOX code in representing tissue identity is largely retained in cancer cell lineages. By applying the HOX code to transcriptomic data from over 114 cancer types, we identified lineage-plastic cancers, such as PRSCC, SCLC, and AML, that exhibit distinct HOX codes compared to their non-plastic counterparts. These findings underscore the potential of HOX code analysis in determining the plasticity status of cancer cell lineages—a critical treatment resistance mechanism—which could improve prognosis and inform clinical decisions.

The HOX code, defined by the spatially and temporally coordinated expression of Homeobox (HOX) genes, serves as a critical determinant of embryonic patterning and cell lineage identity during normal development [30]. However, early exploration of the HOX code across pan-cancer types was limited due to insufficient sample sizes. Through comprehensive bioinformatic analysis of multi-omics data—including expression profiles of 39 *HOX* genes from over 80,000 RNA sequencing samples across 114 cancer types—we demonstrated that HOX codes effectively represent the lineages of cancer cells. Our research represents a significant advancement in translating developmental biology concepts into cancer research applications. While HOX codes have long been recognized for their role in embryonic development and tissue specification, our study expands this concept into cancer biology, providing novel insights into how these developmentally regulated genes contribute to cancer lineage plasticity. By establishing the HOX code as a universal indicator of cancer lineage identity, our work addresses a major limitation in current cancer classification approaches. Unlike canonical cancer lineage markers, which are often loosely defined and vary considerably across different cancer types, HOX code analysis offers a standardized, genome-wide approach to assessing lineage identity regardless of cancer origin. This universality enables more consistent classification of cancer subtypes and more reliable identification of lineage-plastic states across diverse malignancies.

Using this newly developed HOX code, we identified lineage-plastic tumor subtypes in prostate cancer, lung cancer, and AML, which exhibit altered HOX codes compared to their non-plastic counterparts. More notably, our analysis revealed significantly elevated *RUNX1T1* expression across all three lineage-plastic cancer types, which was further validated through both bulk and single-cell RNA sequencing data derived from preclinical and clinical samples. Together, our findings provide a novel strategy for characterizing lineage plasticity in pan-cancer cells through HOX code analysis and further suggest ectopic *RUNX1T1* expression as a pan-cancer marker and critical mediator of lineage plasticity. This addresses a major gap in utilizing lineage plasticity as a predictive biomarker for targeted therapy resistance. Moreover, our findings have substantial clinical implications, particularly in the context of treatment resistance. Lineage plasticity has emerged as a key mechanism underlying resistance to targeted therapies and immunotherapies. By enabling detection and monitoring of lineage plasticity through HOX code analysis, our study offers a potential strategy for improving patient prognosis and guiding therapeutic decision-making. Clinicians could leverage HOX code signatures to identify patients at risk of developing treatment resistance due to lineage plasticity and adjust treatment regimens accordingly.

Collectively, our study establishes HOX code analysis as a powerful approach for detecting cancer lineage plasticity, offering both biological insights into cancer evolution and practical applications in clinical oncology. This work lays the foundation for future studies to explore the mechanistic relationship between *HOX* gene regulation and lineage plasticity in cancer.

## Methods

### Data Sources and Preprocessing

We analyzed bulk RNA sequencing (RNA-seq) datasets from sources including: the pan-cancer cell line transcriptome atlas (PCTA, https://pcatools.shinyapps.io/PCTA_app/), the OHSU AML cohort from cBioportal (https://www.cbioportal.org/), lung cancer patient data from Gene Expression Omnibus (GEO, https://www.ncbi.nlm.nih.gov/geo/). The PCTA dataset comprises

TPM (Transcripts Per Million) and read count values for human GRCh38 genes across multiple cancer cell lines. Metadata for PCTA samples, including cell line identifiers, cancer types, and cancer anatomic sites, were retrieved from a previous publication [37]. For single-cell RNA-seq validation, we utilized two prostate cancer datasets: the Human Prostate Single-Cell Atlas (HuPSA) and the Mouse Prostate Single-Cell Atlas (MoPSA) retrieved from previous publication. HUPSA was filtered to exclude normal and normal-adjacent cells, retaining only “PRAD_AR+_1” and “PRSCC” cell types representing the lineage non-plastic and lineage plastic prostate cancer cells, similarly, MOPSA was subset to “PRAD_1” and “PRSCC” cells from the GEM (Genetically Engineered Mouse) model. Both datasets underwent UMAP dimensionality reduction [38]. Additionally, bulk RNA-seq data from the Proatlas provided TPM values for prostate cancer samples, which were subset to *RUNX1T1* expression for further analysis [38].

### HOX Code Correlation Analysis

Median *HOX* expression per cell line was calculated, forming a matrix that was hierarchically clustered within cancer type groups using Euclidean distance and Ward.D2 linkage. The resulting correlation matrix was visualized as a heatmap with diagonal preservation using ComplexHeatmap, split by cancer type and annotated with cancer anatomic sites. Similarly, median *HOX* expression per cancer type was calculated and visualized as an upper triangular heatmap.

### Differential Expression Analysis

Differential expression analysis was performed using DESeq2 on raw count data from PCTA. Target cell lines were grouped as AML non-plastic (MOLM13, MOLM14), AML plastic (HEL, OCIAML3), prostate non-plastic (22RV1, C42B, PC3), prostate plastic (H660), lung non-plastic (CALU3, H1993), and lung plastic (DMS454, H661, H82). Counts were subset to these groups, prefiltered to retain genes with ≥10 counts in ≥3 samples and analyzed. Results were filtered for upregulated genes using thresholds of log2 fold change (log2FC) > 3 and adjusted p-value (padj) < 0.01, visualized with volcano plots using EnhancedVolcano, highlighting *RUNX1T1*.

### Statistical Analysis and Visualization

All statistical analyses used R (v4.2.1). Correlation heatmaps employed ComplexHeatmap with colorRamp2 for color gradients. Boxplots and violin plots used ggplot2. Venn diagrams for overlapping genes were created with VennDiagram. Plots were saved as PDFs with specified dimensions.

## Code and data availability

All data used in this study is listed in the method section. All code necessary to reproduce every figure in this study is published in GitHub (@JYY2001).

## Acknowledgments

This work was supported or partially supported by: National Cancer Institute/National Institutes of Health: R00CA218885, R37CA258730, R01CA288820, R01CA292949 P. Mu; Department of Defense: W81XWH-18-1-0411 and W81XWH21-1-0520 P. Mu; Cancer Prevention Research Institute (CPRIT): RR170050, RP220473, P. Mu, Prostate Cancer Foundation: 17YOUN12 P. Mu, Yale Cancer Center CCSG Pilot Grant. P30CA016359 P. Mu. and I.Y.K.

